# Seasonality effects and field-estimation of colony size in desert ants

**DOI:** 10.1101/2023.06.30.547264

**Authors:** Aziz Subach, Darar Bega, Maya Saar

**Affiliations:** Entomology and Nematology Department, University of Florida, 1881 Natural Area Dr., Gainesville, FL 32608; School of Zoology, Faculty of Life Sciences, Tel Aviv University, Haim Levanon St. 30, Tel Aviv, Israel 6997801

**Author notes:** Correspondence: Maya Saar.

**Keywords:** *Cataglyphis*, colony size, desert ants, field tools, seasonality

## Abstract

The colony level of eusocial insects is considered the reproductive unit on which natural selection operates. Therefore, seasonal demographic movements and estimations of colony size are crucial variables. Excavating colonies of ants to extract their size is daunting, unhealthy to the surrounding environment, and it may prevent long-term research, including testing seasonal effects on colony size. Previous capture-recapture methods that avoid excavating colonies have been proven inefficient when sampling mostly underground dwellers as ants. To address this issue, we offer a simple method to estimate the colony size of desert ants (*Cataglyphis niger*) in a field setting- based on a field experiment, a literature review, and four laboratory experiments. First, we find that between 10-15% of the colony size are outgoing foragers. Second, we find seasonal effects on colony size and foraging activity: colony size varies and is larger in winter than in summer, and in contrast - the proportion of foragers out of colony size is higher in summer than in winter. This suggests that the energetic requirements of the colonies are higher in summer than in winter. Based on uniquely large sample size, our proposed field method may be useful for other co-occurring *Cataglyphis* species. Moreover, extracting ant colony sizes and evaluating ant biomass is advantageous for future studies to evaluate the carrying capacity of semi sand-dunes habitats.

## Background

Organisms change with seasons; therefore, this should be considered when studying them. Seasonality influences the migration activity, feeding, and mating behavior in many insect species [1]. In the case of social insects, the colony level should be considered when studying seasonal cycles [2,3]. Wilson [4] used the term “sociogenesis” to refer to the birth and long-term changes in colonies of social insects. Sociogenesis includes changes such as worker size variability [5,6], workers longevity [7], or the colony production of sexual and reproduction [8,9]. Colony size is a prominent variable in sociogenesis because of its great impact on the colony. For instance, in many ant species and other social insects, colony size increases with age (under habitat resource constraints). In older and larger colonies, new castes may appear, or existing castes become more differentiated. i.e., specialization increases with colony size [10,11]. When studying sociogenesis, colony size also appears crucial when measuring other variables: Egg laying rate, alates production, and more variables must be studied in relation to colony size [2]. It is advantageous to census ant colony size in a few annual time points, as it is much influenced by seasonality; in the fire ant *Solenopsis invicta*, colony size changes throughout the year, while colony size peaks in December (winter) and descends to the lowest size in July (summer; [3]). When following marked colonies through seasons, this phenomenon is often called “seasonal polydomy”; in the Argentine ant, *Linepithema humile* colonies disperse in summer, while being more aggregated in all other seasons [12]. *Myrmica punctiventris* split their colony size into smaller colonies in Spring and by the end of summer return to form a higher colony size [13,14]. In some species of ants, regulation of polymorphism in castes (both workers and reproductives) is constraint to season [15,16]. Lastly, in leaf-cutter ants, nest entrance number and size vary with seasons [17].

A well-known method to estimate population size in the wild is the capture-recapture method [18]. In this method, individuals from a population are captured and marked, and then released back to the wild to mix with individuals from the same population. After a given period, which varies among species and habitats, marked and unmarked individuals are recaptured. The simplest and most common way to estimate the population size is using the Peterson-Lincoln index (reviewed in [19], and in ants; [20]):

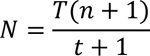

Where *N* is the population size, *T* is the number of marked individuals that were released for the first time, *n* is the number of recaptured individuals (marked and unmarked), and *t* is the number of recaptured marked individuals. There are a few essential assumptions that need to be fulfilled for this method to work: (1) animals are unharmed by marking and the markings persist through the time needed for the population assessment; (2) every individual in the population has an equal probability of being captured and recaptured, regardless of being marked or not and regardless of age, sex, and other traits; (3) the marked individuals can mix back within their original population; (4) sampling time must be lower than the overall time taken for population assessment; (5) the population is closed, meaning there is no immigration or emigration; and (6) there are no births and deaths during the time the population is assessed [19,20].

Some studies on wild ant colonies have used the capture-recapture method in order to estimate colony size [21–25]. However, at least two assumptions of this method (2 and 3) are not met when assessing the size of ant colonies, as stressed in other studies [20,26,27]. Moreover, Erickson [27] demonstrated a steep underestimation of colony size (83-92%) in *Pogonomyrmex californicus* using capture-recapture methods, and [28] showed that ant grooming tends to eliminate marks used. Regarding the second assumption, only up to ∼30% of the colony population are foragers or scouts (Table 1) that can be found outside the nest, while most colony members reside underground, including reproductive members. Therefore, the probability of capturing individuals on site is not equal for all individuals. For the same reason, the third assumption is also not met. Marked individuals will not necessarily mix equally in their original population before recapture (but see: [29]). For instance, while using capture-recapture methods followed by excavation, marked foragers were found only in the upper chambers near the surface in three species of *Pogonomyrmex* [30].

**Table 1:**
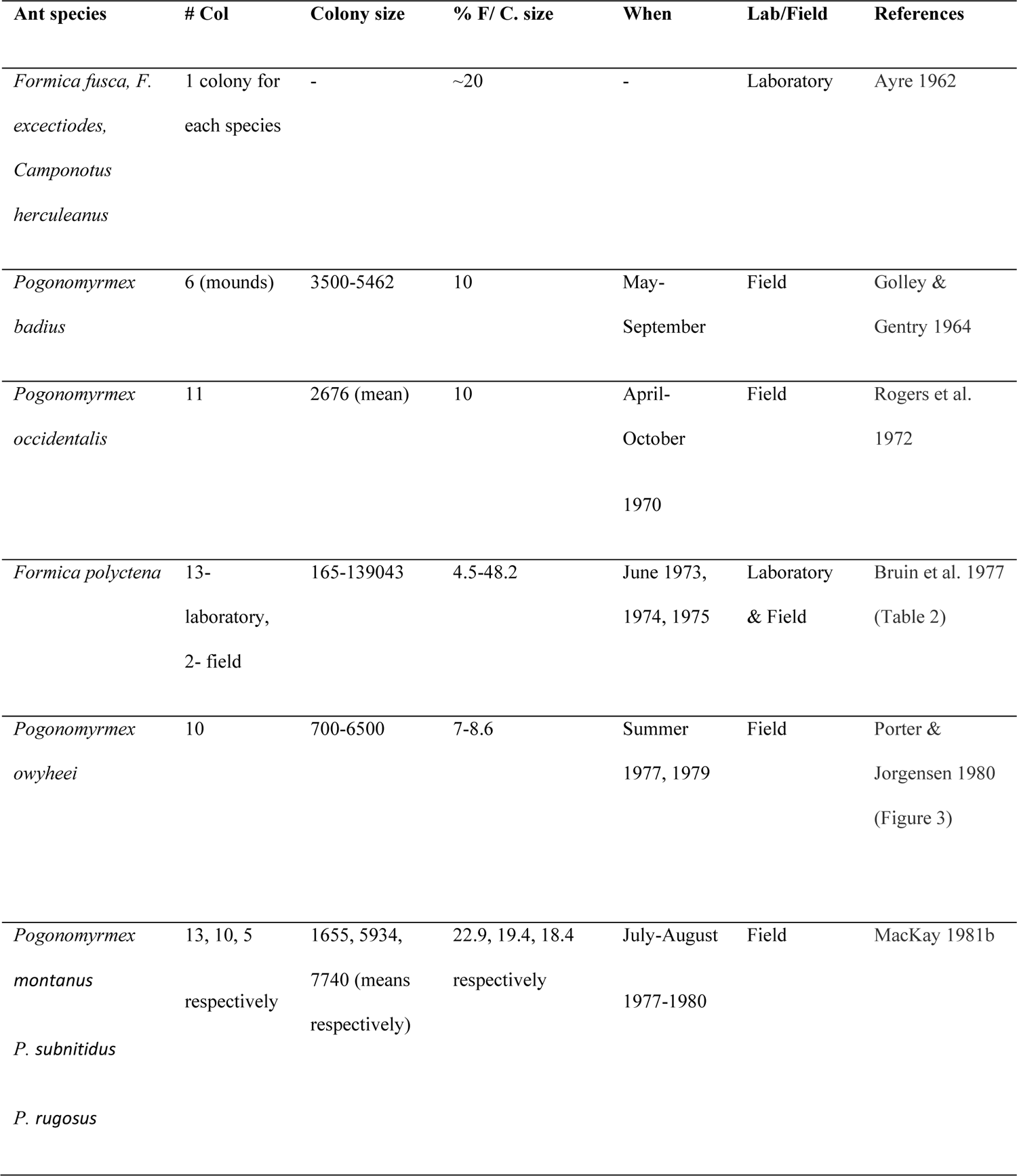
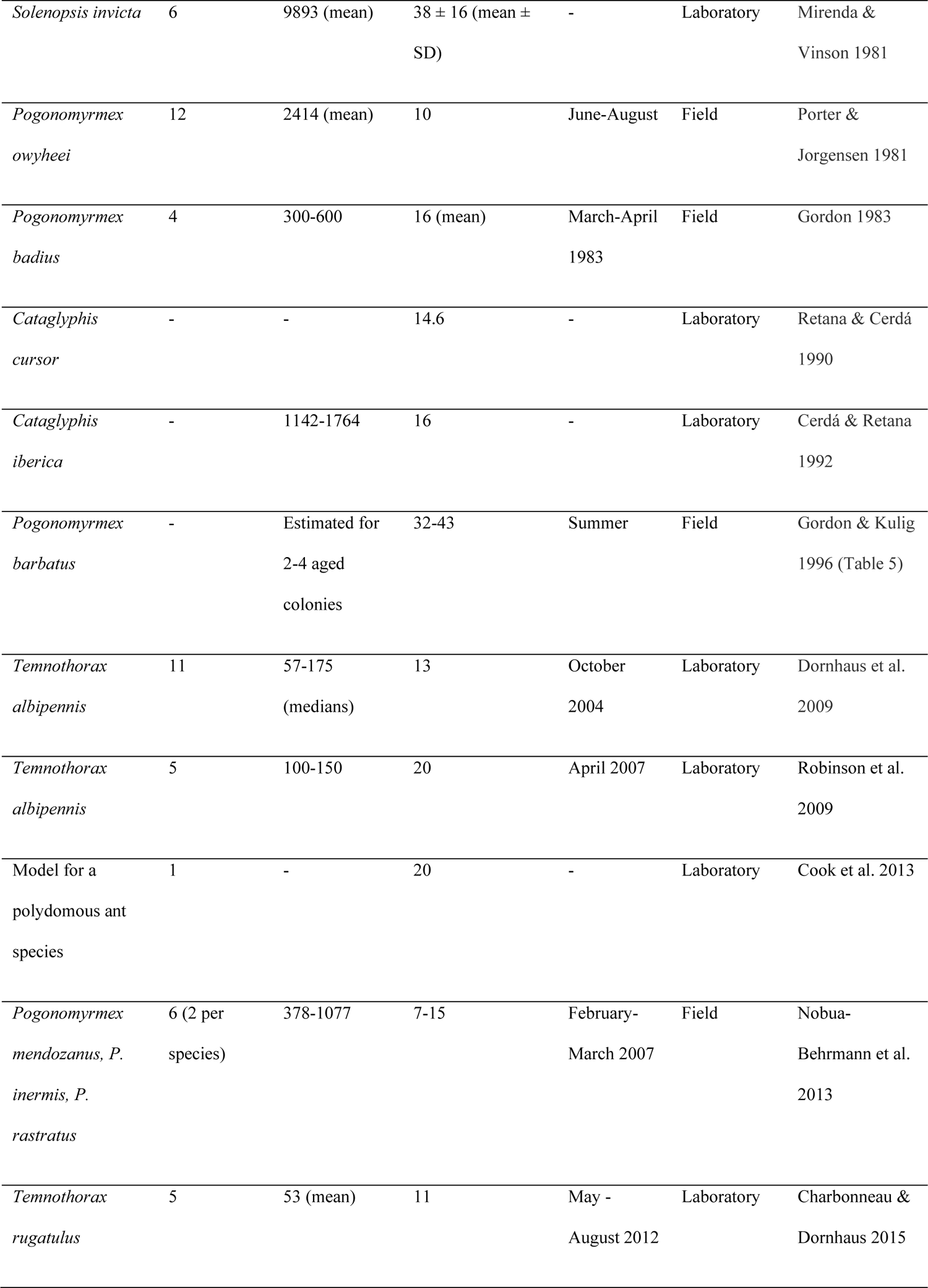
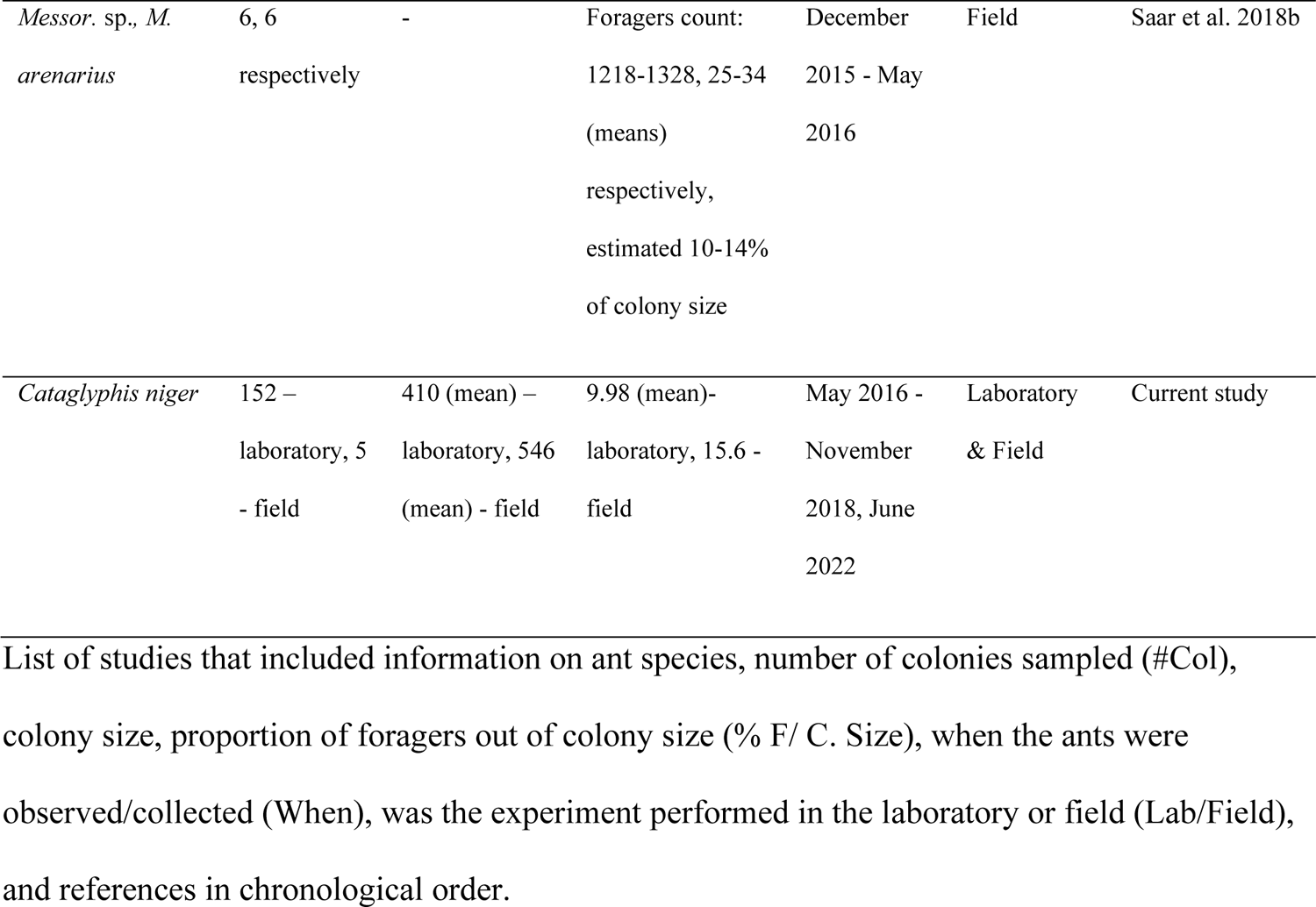
Literature review on colony size and proportion of foragers out of colony size, in various ant species.

The *Cataglyphis* genus comprises ∼100 species that dwell in the Palearctic desert belt and in the Mediterranean basin [31,32]. Past studies that excavated colonies of *C. niger* in Israel, reported on a colony size between 31-3000 individuals [33–36]. Some reported averages such as Leniaud et al. [35]: 730.02 ± 243.34, (mean ± 1 STDV), but all the studies above reported relatively low sample sizes (1-9 colonies) and were randomly excavated across the year. The current study used the highest sample size known of *C. niger* colonies and reported it throughout the year.

Our uniquely large sample size in the present study implies that our proposed field method may be useful for further study of *C. niger*, and possibly for studying other co-occurring *Cataglyphis* species, specifically as evidence accumulated recently that *C. niger* may be a species complex in Tel-Baruch sand dunes (*C. niger*, *C. savignyi*, and *C. drusus* in: [37,38]). However, for coherence and for the lay reader, we leave species classification for further study and refrain from referring to *C. niger* as a *C. niger* complex, until its taxonomic issues are officially resolved.

One goal of the current study was to offer a thorough field experiment to estimate the colony size of *Cataglyphis niger* and possibly other congeneric species. Therefore, we sought to establish a method to estimate colony size in the wild without the need to excavate and desiccate the colony. We relied on a field experiment, a literature review that demonstrated that up to ∼30% of the total colony size were foragers in various ant species, and on data from four laboratory experiments [39–41]. A second goal was to test whether seasonality affects colony size and foraging activity, which should be considered when assessing colony size. We rely on collection of over 200 colonies of *C. niger* in one semi sand-dunes habitat, throughout different seasons, and in ∼2.5 years of research.

## Results

### Descriptive statistics

#### Field experiment

The average colony size for five tested colonies of *C. niger* in the field was 546.2 ± 62.6% (mean ± 1 SE). The proportion of foragers out of colony size for five tested colonies was averaged at 15.567 ± 0.44% (mean ± 1 SE). The distribution of outgoing foragers for 5 control colonies and 5 test colonies is bimodal with two main time points of activity: 07:00-09:00, and 12:00-14:00, but the distribution of the test colonies is more acute compared to the control (Figures 2a, b).

**Figure 1:**
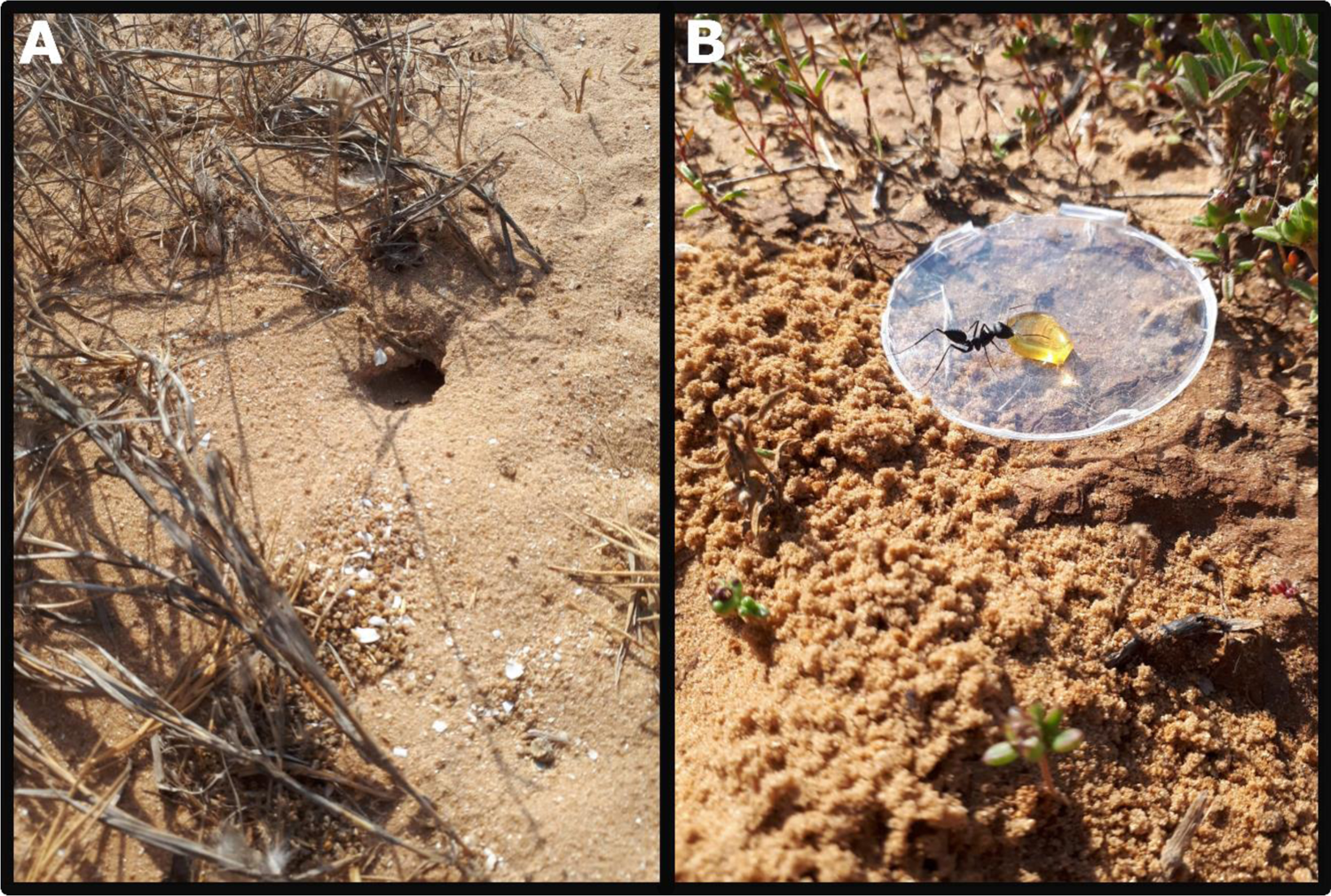
(A) Typical nest entrance of *C. niger* in Tel-Baruch sand dunes. The entrance is at a slight angle from the ground, enabling a rear blind spot for an experimenter to stand and observe ant behavior. (B) *C. niger* feeding on a drop of honey (5g) on a 9 cm Petri dish, cut in its margins to allow easy access, in Tel-Baruch semi sand-dunes. Photos: Bar Avidov.

**Figure 2:**
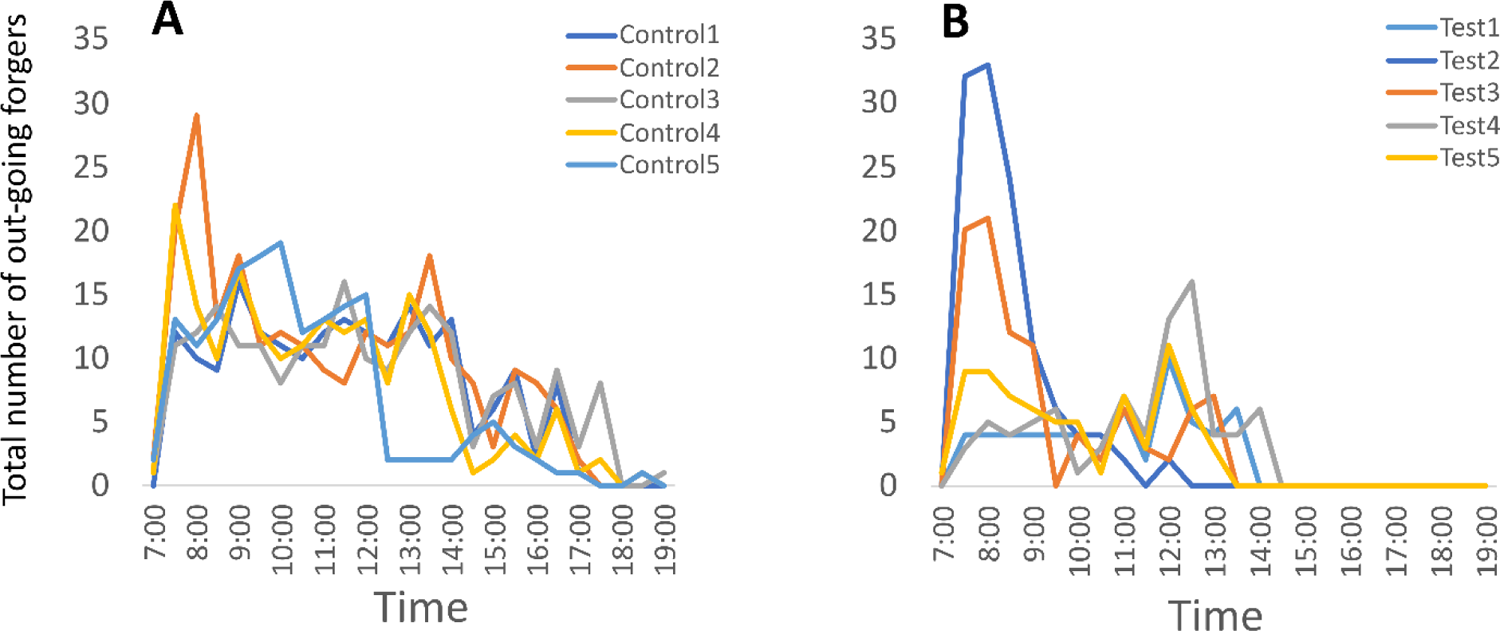
The distribution of outgoing foragers during the field experiment for: (A) five control colonies; and (B) five test colonies, during the 12h period (07:00-19:00) of the experiment.

*Effects of seasonality on colony size and proportion of foragers out of colony size:*

#### Proportion of foragers out of the total colony size

152 colonies were analyzed. 9.98 ± 0.51% (mean ± 1 SE) of the colony foragers left the nest to search the mazes. In summer, the proportion of foragers was 12.07 ± 0.75% (mean ± 1 SE) and in winter −7.67 ± 0.53%.

#### Colony size

The average colony size in the 222 colonies analyzed was 409.75 ± 31.58 (mean ± 1 SE). The smallest colony contained 75 individuals, while the largest colony contained 3218 individuals. In summer we collected 85 colonies with a colony size of 168.96 ± 8.06 (mean ± 1 SE) and in winter we collected 137 colonies with a colony size of 559.14 ± 46.62 (mean ± 1 SE). Both variables-colony size and proportion of foragers out of colony size, are distributed in an asymmetric left distribution (Figures S2a, b).

### Statistical analysis

1. *One-way ANOVA analysis*

a. *Proportion of foragers out of total colony size:* In summer, colonies of *C. niger* released more foragers to search the mazes, relative to colony size, compared to winter (F (1, 150) = 19.58, p < 0.001, Figure 3a).
b. *Colony size:* the colony size of *C. niger* was larger in winter than in summer (F (1, 220) = 96.47, p < 0.001, Figure 3b).
2. *Chi-Square tests*

a. The proportion of foragers out of colony size in laboratory and field colonies did not differ on the same dates that they were tested, three years apart (Table S2, S3; X^2^ (1, 4) = 1.944, P = 0.58).
b. The total number of outgoing foragers did not differ between test and control colonies in the field experiment (Table S4; X^2^ (1, 5) = 8.267, P = 0.08).

## Discussion

Our study highlights life history traits of *C. niger*: Colony size was larger in winter than in summer. In contrast, colonies sent out more foragers to search the maze in summer than in winter. Importantly, we suggest a method that can contribute to field biology studies by enabling the evaluation of the colony size of desert ants in their natural habitat without desiccation. Our Method is based on our field experiment, four laboratory experiments, and literature review that support it. The proportion of foragers out of total colony size was between ∼10-15.6% in our laboratory and field experiments. These proportions generally correspond to other studies in ants, and specifically in the *Cataglyphis* genus ([42,43]; and see Table 1).

**Figure 3:**
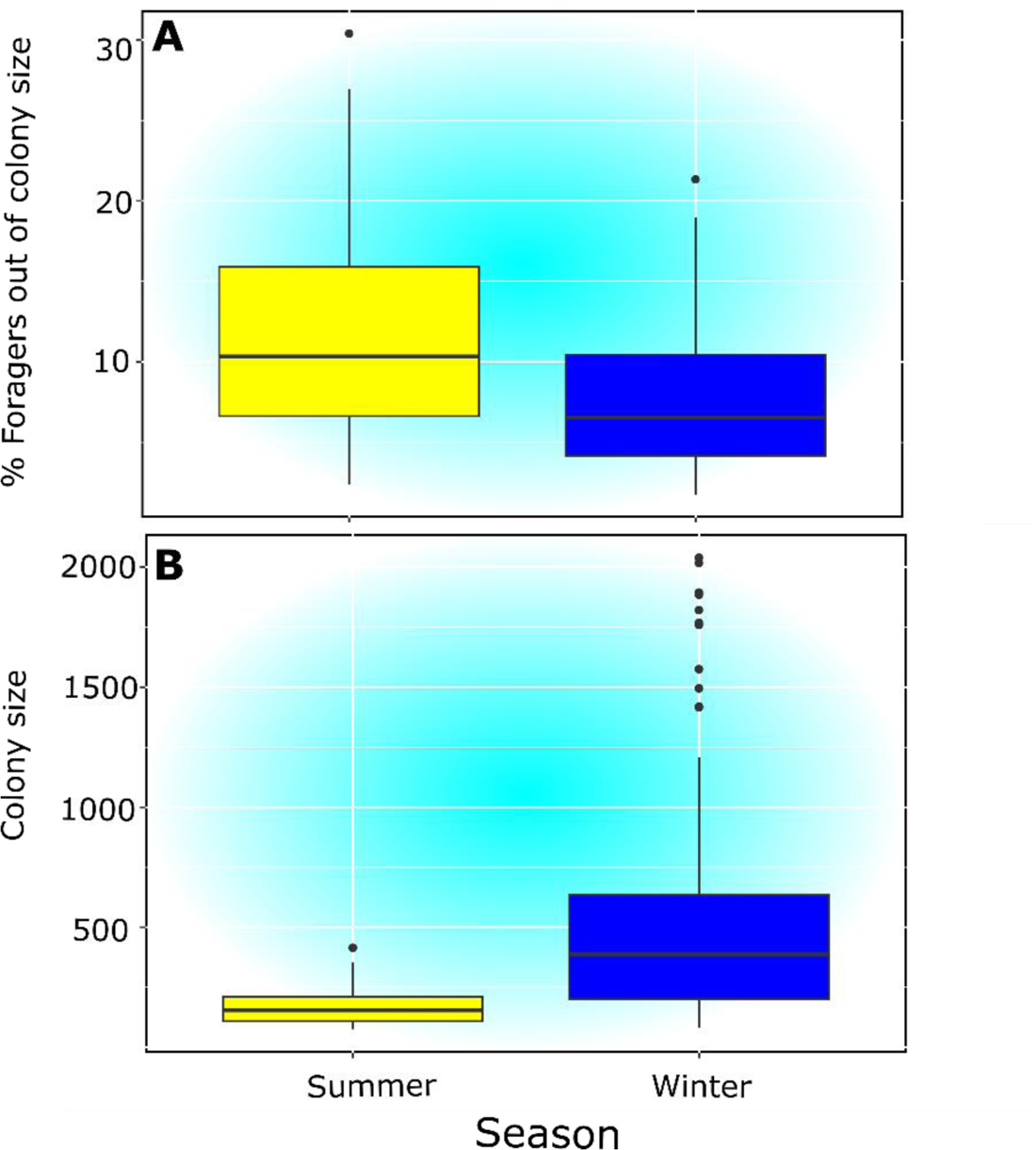
(A) In summer, *C. niger* colonies significantly released more foragers to search the mazes, relative to colony size. (B) Colony size of *C. niger* was significantly larger in winter, than in summer.

A review of 18 past studies depicts colony sizes of ant species from various subfamilies, the proportion of foragers out of the colony size, when in the year it was observed and was it in laboratory or field settings (Table 1). The review demonstrates in a wide species array, that only up to 15 colonies per species were sampled to infer proportion of foragers out of colony size. The sample size in the current study is unparallel to past studies (N=152). In addition, while we carefully counted and reported colony sizes - in some studies the sample size was not reported, or the colony sizes and proportion of foragers were not straight forward reported, and they had to be extracted from the data sets by present authors. This review also demonstrates the gap in exhaustion of time periods examined in our data set and throughout seasons; May 2016-November 2018. In comparison, other studies demonstrated shorter time periods; usually up to 7 months in one year or an examination of a single season in one or a few years. Our one-of-a-kind large and annual sample size may imply that this method could be applied to other *Cataglyphis* species, particularly as currently *C. niger* is undergoing new species classification and may be considered a species complex, encompassing *C. savignyi* and *C. drusus* [37,38]. Moreover, previous studies in congeneric species demonstrated similar proportion of foragers out of colony size: 14.6% in *C. cursor*, and 16% in *C. iberica* [42,43].

Our proposed field method evades the need to excavate entire colonies of *C. niger.* Excavating ant colonies is laborious and destructive. Additionally, as scavenger ants such as *Cataglyphis* demonstrate tight relationships with their environment, excavating colonies may also disrupt the environment that *Cataglyphis* occur in, and preclude further study on desert ants or the environment in general. Many past field experiments using *Cataglyphis* ants that did not estimate colony size were nevertheless crucial for understanding the natural behavior of these ants (for instance foraging behavior [44,45]; Adult transport [46]; Interspecific competition [47]). A prominent disadvantage while testing natural behavior in the field is the lack of colony size.

This variable is important to explain variation in behavior between colonies. For instance, was a colony more efficient in foraging compared to intraspecific colonies, or was it just larger in colony size, meaning it contained more worker force participating in foraging? Was a colony stronger in competing with another species compared to other intraspecific colonies, or was it, once more, just larger in size and therefore more successful in driving the competing species away? One possibility is to excavate the colonies when the experiments are over in order to obtain the colony size (as done for example in [46,48,49]). However, this method might miss the colony size at the time point of the experiments. In short, on one hand, one may want to test the natural behavior of desert ants, but in order to do so fully, one may need their colony size. On the other hand, obtaining colony size by excavating disrupts the natural behavior of desert ants and the environment. If we exclude the capture-recapture method because of inefficiency in ant studies (see Introduction), our suggested method uses a middle way: for example, one can document the number of foragers during their activity in one day, estimate the colony size, and then initiate the desired field experiments. If long-term field experiments are desired, one can estimate colony size in a few time points of these experiments. Ultimately, our proposed method provides a relevant, up-to-date colony size for field experiments. For an improved evaluation of colony size throughout the year, we suggest a further experiment to our field experiment: performing it twice; once in the winter and once in the summer, as our laboratory data show that colony size of *C. niger* differs between these seasons.

Physical features of social insect colonies have been used before to assess colony size without disturbing colony life; physical nest size in termites increases linearly with population size; thus, nest size may be used as an indication of population size ([50,51]; also reviewed in [51]). In the fire ant *Solenopsis invicta*, colony size can be determined in the field by the physical nest mound size, which increases with total ant biomass and varies during the year’s seasons according to this biomass [3,52,53]. Our purposed method adds on to these known methods, not with an external physical feature of the colony but rather more directly, by counting the number of foragers during one day of observations and estimating colony size accordingly. Similarly, in *Myrmica* sp., counting foragers climbing on a stick inserted through the colony entrance has been shown to be a good proxy for colony size [54].

Scavenger ants contribute to nutrient recycling in their habitat soil, and the greater the species diversity, the greater the contribution [55,56]. The majority of the species of the ant genus *Cataglyphis* feed often on dead arthropods and function as scavengers in their habitat [57–60]. They are evidently efficient: *C. bicolor* foragers have a life expectancy of only six days due to predation pressure, but during that time they collect food that weighs 15 to 20 times their body weight [57,61].

Carrying capacity translates to the maximum population size that may be sustained in a given habitat [62–64]. Extracting ant colony sizes on a vast territory and evaluating ant biomass is advantageous for future studies to evaluate the carrying capacity of the habitat. This may be done by sampling the habitat with grids (as in [65]) and sampling all *C. niger*’s colony sizes on grids.

Lahav et al. [66]weighed *C. niger* workers from Tel-Aviv, Israel using an analytical balance to an accuracy of 0.1 mg. On average, workers weighed 36.5 + −0.9 mg (mean+-SE, *N*=92). Future research may concentrate on evaluating colony size through our proposed method, and with the information on workers’ weight from Lahav [66], one can obtain a rough estimation of the colony biomass. Doing so for each colony sampled in grids would estimate the habitat carrying capacity for the biomass of these ants. A good example for studies that had the necessity to evaluate carrying capacity of a habitat-used careful census of gerbil species occurrences and densities and as a result-found differences in spatial and temporal habitat use [67–70]. Without knowing the average weight of each species of gerbil and without knowing the carrying capacity of their habitat - authors could not have made their conclusions.

We have found that *C. niger*’s colony size is larger in winter than in summer, in Tel-Baruch semi sand-dunes habitat (Figure 3b). Energetic balance of ant colonies is often affected by seasonality. Fire ants (*Solenopsis invicta*) and harvester ants (*Pogonomyrmex badius*) store body fat in the fall to be able to survive and overwinter and produce reproductive offspring in the following spring [3,71,72]. But storing body fat can additionally come in the form of adding more of the ‘repletes’ caste to the colony, and therefore result in an increase of colony size [73].

In summer, colonies released more foragers relative to colony size to search the mazes, compared to in winter (Figure 3a). We speculate that in the wild, colonies may send out more foragers to compensate for smaller colony size compared to in winter. Additionally, in the Tel-Aviv area, the mean temperature in summer is 27° C (July), while in winter −14° C (January). *Cataglyphis* ants are scavengers that can cope well with heat conditions [74,75]. Therefore, perhaps sending more workers to forage in summer is worthwhile, as heat-struck arthropods are more abundant compared to wintertime. In the field, and in resemblance to *C. niger*, *C. bombycina*’s annual activity and that of two other species of *Ocymyrmex* desert ants were tied to the fluctuations of the surface temperatures, peaking during the hottest hours of the day-close to lethal limits, and in a bimodal distribution in summer, but not in winter [45]. The activity of *C. niger* also peaked in a bimodal distribution during our field experiment - at the beginning of the day (07:00-09:00), and for most colonies, between 12:00-14:00-within the hottest hours of the day in its habitat, and similarly to *C. floricola* in July (see Figure 4b in [76]). As we did not execute our field experiment in winter, we do not know if the distribution of outgoing foragers of *C. niger* could have changed; this remains an endeavor for further research. These two outbursts of foragers from the nest entrance, that are shown in the bimodal distribution in summer, may have been elicited by mandibular gland secretions [77,78], in response to an appropriate surface temperature near the nest or solar cues perceived [45,76]. Interestingly, test colonies of *C. niger* demonstrated a more acute bimodal distribution and a shorter activity time during the day-compared to control colonies. This may have been a response to the continuous removal of foragers by the experimenter. However, as numbers of outgoing foragers stayed stable between test and control colonies, we speculate that tested colonies did not stop the flow of foragers as a response to the removal compared to control (Table S4 and Figures 2a, b).

We stress, despite a small sample size, that we did not find a significant difference in the proportion of foragers out of colony size-between field and laboratory colonies examined. We therefore, incline to suggest the integrity and complimentary aspects of the current study (Tables S2, S3).

## Conclusions

We demonstrate that the colony size and foraging activity of *C. niger* is tied to seasonality. In addition, we provide a straightforward and efficient method to estimate the colony size of *C. niger* in a field setting. Our uniquely large data set implies that this technique may be useful for estimating the colony sizes of co-occurring *Cataglyphis* species. Importantly, this technique may lay the ground for our understanding of the capacity of the habitat to carry such influential scavenger ants.

## Methods

Field experiment to estimate the colony size of wild *C. niger*:

### Preliminary

We observed the activity of 5 colonies throughout one day to determine the beginning and end of the foraging activity. *C. niger* is diurnal, and we therefore arrived before sunrise and left after sunset.

### Experiment

The experiment took place over five days during June 2021: June 20, June 21, June 22, June 28, and June 29. For each date, one experimental colony was marked, and all five were 500-800 meters apart. 30 meters next to each experimental colony, we marked a control colony. All colonies (experimental and control) were given a 9 cm Petri dish with pre-weighed 5g of honey, 30 cm from the nest entrance, 24 h before the experiment and were left for 12 hours for habituation of the ants to the experimental protocol. On the day of the experiment, each experimental colony was treated with the following protocol:

A 9 cm Petri dish with 5g of honey was inserted into the sand, 30 cm from the nest entrance until its edges were equal to the ground level for easy access for ants. An experimenter established an observation point 5 m behind the nest entrance and used a 10X42 Nikon Prostaff P3 binocular to document the activity of ants. In many cases in Tel Baruch sand dunes, the colony entrance has an angle to the ground (Figure 1a); thus, standing behind the entrance would make the ants almost blind to the experimenter’s presence. The experiment lasted between 07:00 and 19:00 (12 hours), encompassing all hours that ants were expected to be active. From 07:00 am, every 30 min (following [45]), foragers feeding in the Petri dish were collected into a plastic ventilated container (20X15 cm) by the experimenter and were kept there for the entire experiment. The experimenter noticed that only foragers that emerged from the tested nest entrance accessed the Petri dish. In a few instances, non-nestmates tried to access the Petri dish and were aggressively driven away by the nestmates. We expected wild ants to approach the honey-filled Petri-dish because we have evidence that *C. niger* prefers honey over other food baits [39,41]. The experimenter waited for the Petri dish to be filled with feeding ants (Figure 1b) and collected them at intervals of 30 min. Ants in Petri-dish concentrate on feeding and therefore are very easy to collect. The experimenter collected these foragers and placed them in a plastic ventilated box (15X20 cm). When periodic collection was over, the experimenter returned to its backstage observation point and waited for another 30 min for the Petri-dish to re-fill with feeding foragers for further collection. The experimenter repeated collection from the Petri dish as needed until foragers stopped emerging from the colony entrance. We suggest that the colony maintains the flow of outgoing foragers even when they are collected (e.g., when foragers do not return to the nest). Laboratory experiments indicated that when foragers are removed, the colony produces new foragers in a similar proportion (∼10% of colony size: [39–41]). Nevertheless, we performed a Chi-square test between tested and control colonies during the field experiment to check if they might differ in the total number of outgoing foragers (see Methods and Results). The morning after the experiment, prior to the beginning of the foragers’ activity, each of the 5 experimental colonies was excavated carefully to the full extent possible and placed in designated plastic ventilated boxes (15×20 cm). In the laboratory, for each colony, ants from the two boxes (foragers and entire colony collected) were counted. We repeated the protocol for the control colonies, but at the end of the day, we released the foragers back to their natural environment, and did not bring them to the laboratory. See Table S1 for a complete list of the experiment, including dates, and how many foragers were counted each 30 min for the experimental and control colonies.

### Effects of seasonality on colony size and proportion of foragers out of colony size

Data were obtained from four experiments performed between May 2016-November 2018 [39–41]. All *C. niger* colonies were excavated in Tel-Baruch semi sand-dunes, North-West Tel-Aviv, Israel (32.1283 N, 34.7867 E; ∼20 m above sea level). For a detailed description of this habitat flora, see Saar et al. [65]. For the purpose of the four experiments, colonies were excavated, brought to the laboratory, and individuals were counted immediately upon arrival to obtain colony size. The date of collection determined the season of collection and thus was recorded.

Season categories were determined using the Israel Space Agency website (https://www.space.gov.il/en), which indicates that June 20 is the longest day in the north hemisphere and thus the beginning of summer, while December 20 is the shortest day in the north hemisphere and the beginning of winter. All colonies were inserted into a two-section cage; one contained the colony permanently and the second contained a maze (Figure S1). All colonies were maintained in a rearing room under a controlled temperature of 28 °C and a photoperiod of 14L:10D, and were tested in the same room during the light photoperiod. The experiments were designed to answer different questions, but all had common features that we used: In all experiments, we used a Petri dish (6 cm) at the end of the maze, mostly containing a food reward (5 g of either honey or squashed crickets). To start the experiments, we opened a sliding door between the colony section and the maze section of the cage, and let the ants roam in the maze in order to search for the food reward. When the first ant arrived at the food reward, we let the ants feed on it for another 10-15 minutes, and then closed the sliding door and counted all the ants that were feeding or still roaming the maze. We defined this variable as “ants searching” (and in the current study we name it “foragers” for simplicity). Following, we returned all foragers to the colony section of the cage for a 30-minute break, before testing them once more in the maze (runs). In the different experiments, we used a different run number (3-18 runs); therefore, the variable “ants searching” was averaged across the run number. 222 colonies were analyzed in total. 85 were excavated in summer and 137 were excavated in winter. Of the 222 colonies, 70 were used for other experiments; therefore, we considered only their collection date (either in winter or in summer) and colony size. We did not have data on the number of ants searching in the maze for these 70 colonies.

## Statistical analysis

We performed

1. *Two one-way ANOVAs* with season (two levels: winter and summer) as the explanatory variable and two response variables: (a) proportion of foragers out of the colony size that we achieved by dividing the variable “ants searching” with colony size (b) colony size. Colony size and proportion of foragers out of colony size variables were both log-transformed due to their abnormal distributions.
2. *Two chi-square tests:* (a) To test the association between laboratory colonies vs. field colonies in the proportions of foragers out of colony size: We compared laboratory colonies that were collected on four out of five dates (20.6, 21.6, 28.6 and 29.6) that field colonies were collected, but three years earlier (Table S3). (b) To test the association between tested colonies vs. control colonies in the total number of outgoing foragers in the field experiment.

## Declarations

### Ethics approval and consent to participate

Not applicable

### Consent for publication

Not applicable

### Availability of data and materials

All data generated or analysed during this study are included in this published article (and its supplementary information files).

## Author’s contributions

AS designed the study and acquired data. MS designed the study, acquired, analyzed, interpreted data, and drafted the manuscript. DB acquired data. All authors contributed to the writing and revising of the manuscript.

## Acknowledgements

We thank Inon Scharf for the useful comments on an early version of the manuscript.

## Authors’ information

Not applicable

