## Supplementary material for "Seasonality effects and field-estimation of colony size in desert ants": Table S1, Table S2, Table S3, Table S4, Figure S1, Figure S2

Number of Tables and Figures: 4 & 2

### Tables

**Table S1**: Field experiment data. We report the number of feeding foragers that were collected every 30 min from the petri dish, from 07:00 to 19:00, in five days during June 2021. T = test colony, C = control colony. Numbers next to either T or C depict the colony ID, and the three rows at the end show: Total number of foragers, the colony size (N), and the proportion of foragers out of colony size for one colony during one day:

| **Hour / date** | **June 20, 2021** | | **June 21, 2021** | | **June 22, 2021** | | **June 28, 2021** | | **June 29, 2021** | |
|---|---|---|---|---|---|---|---|---|---|---|
| **Colonies test/control** | **T1** | **C1** | **T2** | **C2** | **T3** | **C3** | **T4** | **C4** | **T5** | **C5** |
| 7:00 | 0 | 0 | 0 | 2 | 0 | 1 | 0 | 1 | 1 | 2 |
| 7:30 | 4 | 12 | 32 | 20 | 20 | 11 | 3 | 22 | 9 | 13 |
| 8:00 | 4 | 10 | 33 | 29 | 21 | 12 | 5 | 14 | 9 | 11 |
| 8:30 | 4 | 9 | 24 | 13 | 12 | 14 | 4 | 10 | 7 | 13 |
| 9:00 | 4 | 16 | 11 | 18 | 11 | 11 | 5 | 17 | 6 | 17 |
| 9:30 | 4 | 12 | 6 | 11 | 0 | 11 | 6 | 12 | 5 | 18 |
| 10:00 | 4 | 11 | 4 | 12 | 4 | 8 | 1 | 10 | 5 | 19 |
| 10:30 | 2 | 10 | 4 | 11 | 2 | 11 | 3 | 11 | 1 | 12 |
| 11:00 | 6 | 12 | 2 | 9 | 6 | 11 | 7 | 13 | 7 | 13 |
| 11:30 | 2 | 13 | 0 | 8 | 3 | 16 | 4 | 12 | 3 | 14 |
| 12:00 | 10 | 12 | 2 | 12 | 2 | 10 | 13 | 13 | 11 | 15 |
| 12:30 | 5 | 11 | 0 | 11 | 6 | 9 | 16 | 8 | 6 | 2 |
| 13:00 | 4 | 14 | 0 | 12 | 7 | 12 | 4 | 15 | 3 | 2 |
| 13:30 | 6 | 11 | 0 | 18 | 0 | 14 | 4 | 12 | 0 | 2 |
| 14:00 | 0 | 13 | 0 | 10 | 0 | 12 | 6 | 6 | 0 | 2 |
| 14:30 | 0 | 4 | 0 | 8 | 0 | 3 | 0 | 1 | 0 | 4 |
| 15:00 | 0 | 6 | 0 | 3 | 0 | 7 | 0 | 2 | 0 | 5 |
| 15:30 | 0 | 9 | 0 | 9 | 0 | 8 | 0 | 4 | 0 | 3 |
| 16:00 | 0 | 2 | 0 | 8 | 0 | 3 | 0 | 2 | 0 | 2 |
| 16:30 | 0 | 8 | 0 | 6 | 0 | 9 | 0 | 6 | 0 | 1 |
| 17:00 | 0 | 1 | 0 | 2 | 0 | 3 | 0 | 1 | 0 | 1 |
| 17:30 | 0 | 0 | 0 | 0 | 0 | 8 | 0 | 2 | 0 | 0 |
| 18:00 | 0 | 0 | 0 | 0 | 0 | 0 | 0 | 0 | 0 | 0 |
| 18:30 | 0 | 0 | 0 | 1 | 0 | 0 | 0 | 1 | 0 | 1 |
| 19:00 | 0 | 0 | 0 | 0 | 0 | 1 | 0 | 0 | 0 | 0 |
| Total foragers | 59 | 196 | 118 | 233 | 94 | 205 | 81 | 195 | 73 | 172 |
| N colony | 402 | | 731 | | 645 | | 521 | | 432 | |
| % foragers out of N colony | 14.67 | | 16.14 | | 14.57 | | 15.54 | | 16.89 | |

**Table S2:** The proportion of foragers out of colony size in laboratory and field colonies ("col.") did not differ on the same dates that they were tested, three years apart (2018 and 2021) according to chi-square analysis. For each cell we report: the proportion of foragers out of colony size, (the expected cell totals), and [the chi-square statistic for each cell]:

| | **June-20** | **June-21** | **June-28** | **June-29** | **Row totals** |
|---|---|---|---|---|---|
| **Laboratory col.** | 22 (19.46) [0.33] | 20 (18.93) [0.06] | 16 (16.83) [0.04] | 13 (15.78) [0.49] | 71 |
| **Field col.** | 15 (17.54) [0.37] | 16 (17.07) [0.07] | 16 (15.17) [0.05] | 17 (14.22) [0.54] | 64 |
| **Column totals** | 37 | 36 | 32 | 30 | **Total: 135** |

**Table S3**: In four out of five dates, we could compare the proportion of foragers out of colony size in laboratory vs. field colonies ("col."). The table depicts the year and the four dates in June in which the experiments were conducted (both in Bold), the proportion of foragers out of colony size (in Italic), and the N of colonies tested in parentheses. When more than one colony was tested in one day, we averaged the proportion of foragers out of colony sizes:

| | **Year** | **Jun-20** | **Jun-21** | **Jun-28** | **Jun-29** |
|---|---|---|---|---|---|
| **Laboratory col.** | 2018 | *22* (3) | *20* (3) | *16* (3) | *13* (1) |
| **Field col.** | 2021 | *15* (1) | *16* (1) | *16* (1) | *17* (1) |

**Table S4**: The total number of outgoing foragers did not differ between test and control colonies in the field experiment, according to chi-square analysis. For each cell we report: the total number of outgoing foragers, (the expected cell totals), and [the chi-square statistic for each cell]:

| | **Colony 1** | **Colony 2** | **Colony 3** | **Colony 4** | **Colony 5** | **Row totals** |
|---|---|---|---|---|---|---|
| **Test** | 59 (76.00) [3.8] | 118 (104.61) [1.71] | 94 (89.11) [0.27] | 81 (82.26) [0.02] | 73 (73.02) [0.00] | 425 |
| **Control** | 196 (179.00) [1.61] | 233 (246.39) [0.73] | 205 (209.89) [0.11] | 195 (193.74) [0.01] | 172 (171.98) [0.00] | 1001 |
| **Column totals** | 255 | 351 | 299 | 276 | 245 | **Total: 1462** |

### Figures

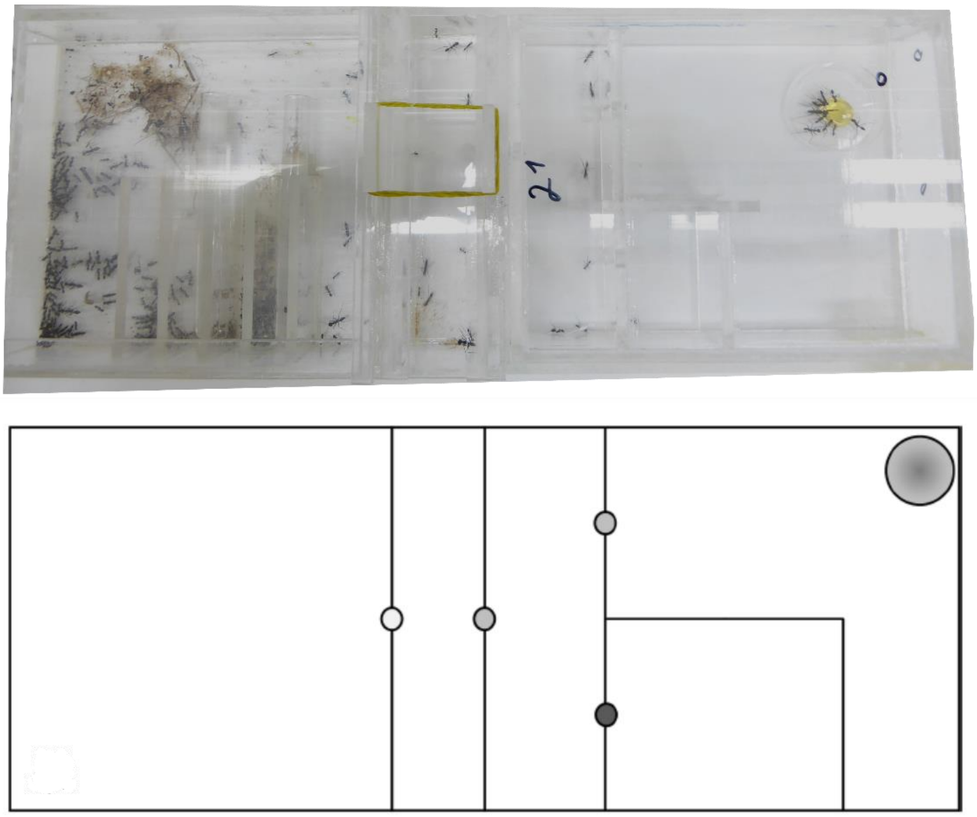

**Figure S1**: Above is a picture of a maze, and below - an illustration. The nest of each colony was housed on the left half of the cage, and two sliding doors were installed between the nest and the maze. When the sliding doors were removed, the ants could have accessed the maze, and the goal was to get to their food reward (largest circle). The grey dots indicate the correct way to reach the food reward, while the black dot indicates a dead end.

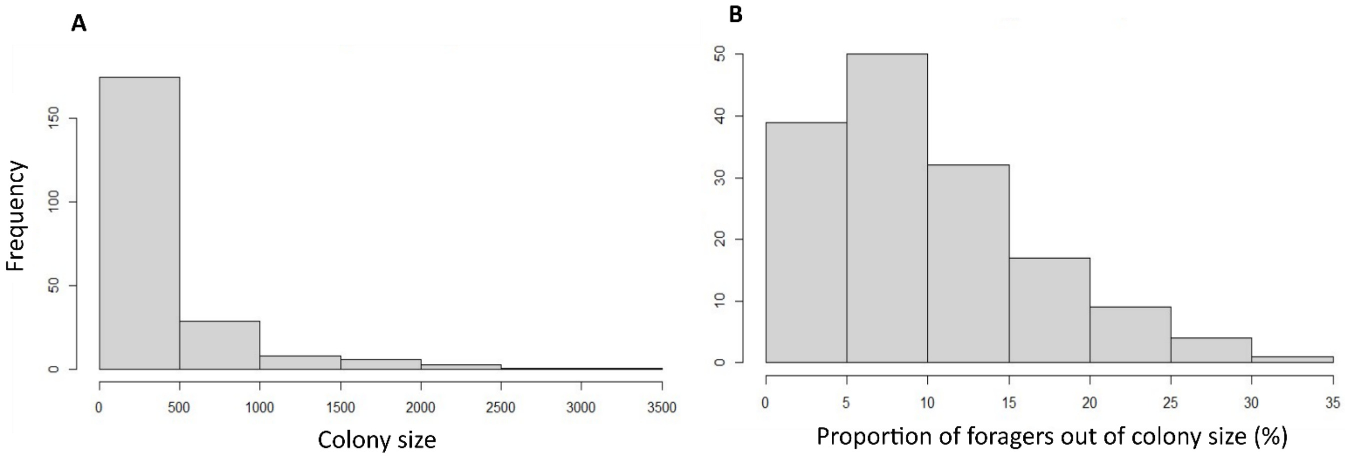


**Figure S2:** a-symmetric left distributions of: (A) Colony size; and (B) proportion of foragers out of colony size.
